# Surrounding Traffic Matters: Increases in Traffic Volume Are Related to Changes in EEG Rhythms in Urban Cyclists

**DOI:** 10.1101/2022.05.27.493782

**Authors:** Daniel Robles, Jonathan W. P. Kuziek, Jingyi Lai, Robin Mazumder, Joanna E. M. Scanlon, Kyle E. Mathewson

**Affiliations:** Department of Psychology, Faculty of Science, University of Alberta

**Author notes:** **Corresponding Author:** Daniel Robles, Department of Psychology, Lab number, P-217 Biological Sciences Building, University of Alberta, Edmonton, Alberta, Canada T6G 2R3.

**Keywords:** Mobile EEG, attention, Cognition, ERPs, real-world EEG

## Abstract

In this study, we used an oddball EEG bicycle paradigm to study how changes in urban environments elicit changes in EEG markers. Participants completed an auditory oddball task while riding in three different cycling lane environments. A low traffic condition where participants rode in a fully separated bike lane alongside a quiet residential street, an intermediate traffic condition where participants rode alongside a busy residential street in a painted lane, and a heavy traffic condition where participants rode alongside fast/heavy traffic on a shared-use path. Relative to the low traffic, heavy traffic was associated with faster reaction time and a trend towards reduced accuracy, and increased N1 amplitude evoked by the standard tones. We attribute this difference in N1 amplitude to different attentional demands evoked by the different traffic conditions. In this fashion, heavy traffic requires greater auditory filtering. Furthermore, we found no differences in P3 amplitude associated with the traffic conditions. We discuss the implications of mobile paradigms to study attention in real-world settings.

## 1. Introduction

### 1.1. Breaking away from the laboratory

From the early days of scientific psychological research until now, most published research has occurred inside research laboratories. This traditional research has prioritized high experimental control during data collection (Bigdely-Shamlo et al., 2016). For instance, to maintain high internal validity, researchers have stressed the importance of controlling for extraneous factors such as light, noise, temperature, etc. (Luck, 2014). When it comes to the study of human attention, techniques such as electroencephalography (EEG) require high standards in experimental control due to its vulnerability to physical movement such as foot-tapping, jaw clenching, eye blinking, etc. For example, it is commonly known that several sources of physical noise such as movement, as well as biological artifacts such as heartbeat and respiration, can cause temporary distortions in EEG signals (Delorme et al., 2007). These artifacts are inherent to EEG and require proper collection techniques (training participants to blink less or in irregular intervals) or by applying correction/rejection techniques (Makeig et al., 2009). Virtually, most of what we know about human brain processes related to attention and cognition comes from laboratory paradigms. Not surprisingly, researchers have previously argued that the dynamic nature of brain processes should be studied under equally dynamic environments, where subjects can walk, reach, navigate, etc. (Gramann et al., 2014; Jungnickel et al., 2018; Ladouce et al., 2017; Julian Elias Reiser et al., 2019).

Fortunately, the evolution of modern technology has brought unprecedented computing power to portable devices such as phones and minicomputers. These advancements have directly enabled researchers’ ability to use EEG in naturalistic actions and environments (Scanlon, Townsend, et al., 2019). A powerful approach known as mobile brain/body imaging (MoBI) was developed to study movement and brain dynamics in three-dimensional environments (Makeig et al., 2009). These brain/body dynamics are captured by collecting data from several sources that include EEG sensors, movement sensors, and the timing of events in the environment. Ultimately, the goal of the MoBI approach is to understand the relationship between active behaviors (moving, touching, pointing, etc.) and brain dynamics associated with these behaviors (Jungnickel et al., 2018). Since artifacts and noise are known to affect EEG quality, mobile approaches rely on data cleaning techniques such as independent component analyses for data cleaning and processing (Gwin et al., 2010).

Previous studies from our research group have focused on the development and use of mobile paradigms. This includes the validation of mini computers such as the Raspberry Pi minicomputer as a valid alternative for EEG data collection (Kuziek et al., 2017), using the Muse headband for stroke prediction in ambulance/ER settings (Wilkinson et al., 2020), and deploying the Muse headband for EEG data collection in rural communities in Malawi (Leal Neto et al., 2021). In several studies, our workgroup used mobile settings to explore the role of several attentional mechanisms during active tasks. For example, we developed and validated an e-skateboard paradigm to study attentional allocation during active skateboarding (Robles et al., 2021). In that study, we manipulated task difficulty by having participants complete an auditory oddball task while riding a Bluetooth-operated e-skateboard around a busy running track. We found that increases in motor difficulty were not associated with changes in the P3 event-related potential (ERP), commonly linked to the reallocation of cognitive resources (De Sanctis et al., 2014). We also found global reductions in alpha power during skateboarding relative to a resting condition. It was concluded that such a change in alpha power could be due to the skateboarding task inducing a state of cortical excitability (Sauseng et al., 2009).

Several studies from our workgroup have used a cycling paradigm to study attention in naturalistic settings. Using an auditory oddball paradigm, authors deployed a cycling EEG paradigm where subjects performed the task while cycling outside and sitting down in a faraday chamber (Scanlon, Townsend, et al., 2019). They found a decrease in alpha power and P300 amplitude outdoors relative to the laboratory condition associated with the reallocation of cognitive resources while cycling. Notably, they found that cycling outdoors was associated with decreases in P2 and increases in N1 amplitude. The N1 is a negative early deflection in the ERP that is generally greater for the attended, as opposed to the ignored, stimuli (Woldorff & Hillyard, 1991). That effect was further replicated in a different study where participants rode a bicycle in a quiet park and next to a busy road (Scanlon et al., 2020). It was found a consistent increase in N1 amplitude alongside the busy road relative to the quiet indoor environment. In both studies, it was concluded that the increase in N1 amplitude could serve as an auditory filtering mechanism to better process the tones in the noisier environment. In a laboratory experiment, Scanlon et al (2019) administered the auditory oddball task while participants listened to various sources of noise (white noise, silence, outdoor sounds). In this study, they found an increase in N1 and a decrease in P2 amplitude when participants listened to a white noise mask and traffic sounds. It was concluded that the N1/P2 modulations could reflect sensory filtering of the background noise, particularly to ecologically valid sounds. By deploying a mobile EEG paradigm outdoors, these cycling studies suggest that different environmental conditions lead to predictable changes in early sensory ERP amplitude as a mechanism of auditory filtering.

Other EEG studies have explored similar indices of attention under naturalistic behaviors such as walking etc. Many of these studies have assessed resource allocation using various electrophysiological indices such as the P3 and EEG frequency spectra such as the alpha and beta band. For instance, Zink et al. (2016) also found that outdoor cycling was associated with a decrease in alpha power and P3 peak amplitude relative to indoors. The authors suggested that an increase in cognitive workload outdoors could be driving the differences in P3 and alpha amplitude. In a dual-task paradigm, Liebherr et al. (2018) showed that both cognitive difficulty and increases in motor demands lead to a more negative P3 amplitude. (Ladouce et al., 2017) concluded that the observed reductions in P3 amplitude could be related to sensory inertial processing (during physical displacement through space) as opposed to the action of walking alone. Interestingly, Storzer et al. (2016) found that, relative to cycling, walking is associated with a greater alpha power decrease. These authors argued that the sensory processing and motor planning associated with walking could lead to the decrease of alpha power observed.

### 1.2. Different infrastructures, different attentional demands?

Cycling has been shown to improve individual physical health, including cardiovascular health (Oja et al., 2011). Furthermore, urban cycling has been shown to improve an individual’s mental well-being (Ma et al., 2021). Given these benefits, it would be important for governments to consider how to encourage cycling for residents. One study suggests that cyclists experience less physiological stress where cycling infrastructure exists (Teixeira et al., 2020). While many distractors can compromise cycling safety, our current study aimed to understand the influence of traffic volume noise in different cycling environments. Inspired by the findings from Scanlon and colleagues suggesting differences in auditory filtering in indoor vs outdoor cycling (2019), and in noisy roadways relative to quiet park areas (2020), the current study aimed to test the effects of traffic volume in different types of bicycle lanes. The current study explores the reallocation of cognitive resources (as measured by the N1/P2 and P300 ERPs and alpha power) during active bicycle riding. Participants completed an auditory oddball task while riding a bicycle in different cycling environments (heavy, intermediate, and low traffic). Using these differences in road types as the experimental conditions, we hypothesized that, relative to low traffic, the heavy traffic riding should be associated with changes in auditory processing related to N1 and P2 amplitude. That is, riding during heavy traffic should increase attentional demands in the bicycle-oddball task. We also hypothesized that relative to a low traffic condition, riding in the heavy traffic condition should lead to decreases in positivity in the P3 ERP and alpha power alike.

## 2. Methods

### 2.1. Participants

A total of 24 individuals from the university community were recruited (mean age = 20.96, age-range = 18 - 27). Participants received an honorarium of 10 dollars/hr. or 2 credits towards Research Participation for courses in the Psychology department. Participants presented no history of physical or neurological disorders. Before the experiment, each participant verbally confirmed that he/she/they were comfortable riding a bicycle. This study was approved by the Internal Research Ethics Board at the University of Alberta (Pro00050069) and participants signed an informed consent form before completing the study.

### 2.2. Materials and procedures

In this experiment, participants rode a Kona Mahuna bicycle (figure 1A). Participants selected the bicycle size (17” or 19”) based on height while the saddle height was adjusted based on the rider’s preference and comfort. The bicycles were equipped with a response button for the auditory oddball task. This button was attached on the right side of the handlebar and adjusted to each participant to ensure that they could respond with their right thumb while riding. This location of the response button allowed for smooth handling, and access to the brake lever while responding to the task without moving the hand out of its natural riding position. In line with the work of Scanlon et al., (2020), the bicycle gear/resistance was kept constant for all participants (second gear in the crankset and half gear in the cassette) for consistency and to maintain the physical activity at sub-aerobic levels throughout trials.

**Figure 1.**
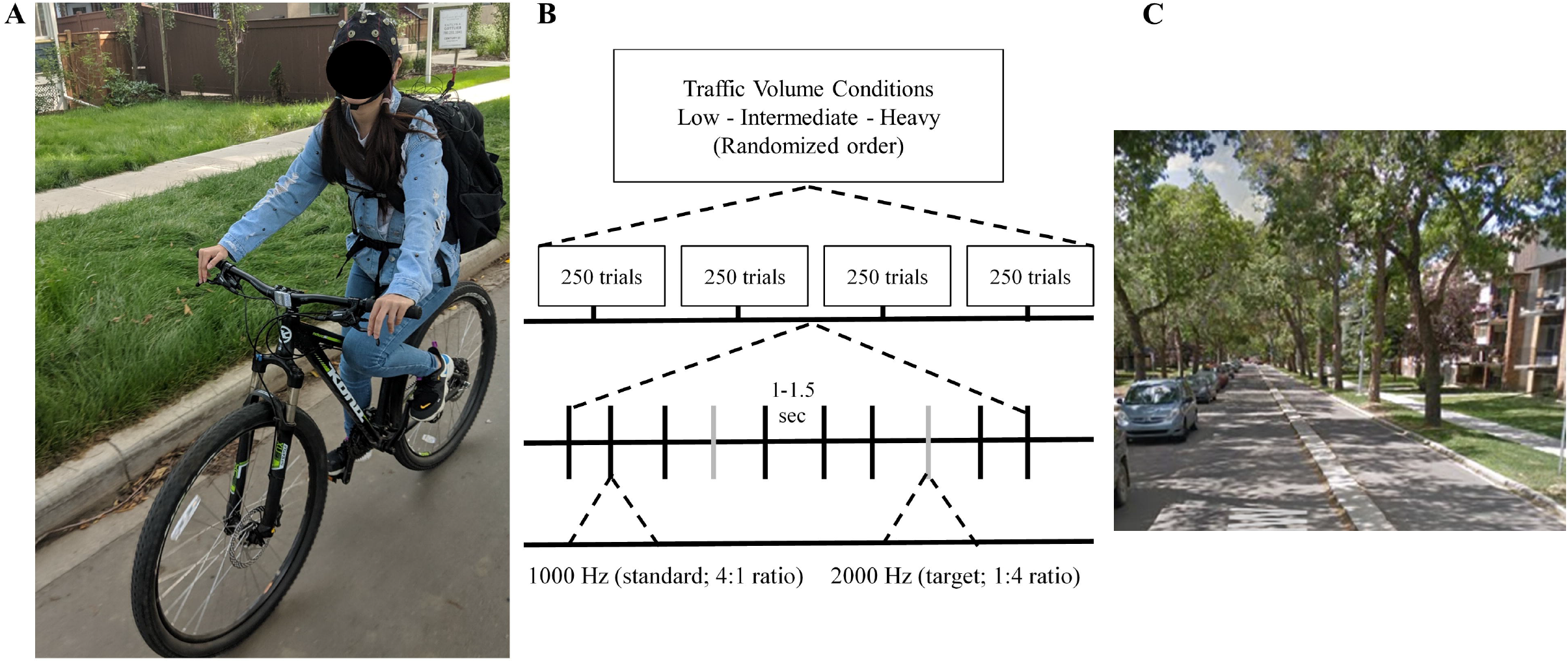
Setup and task. A: Bike EEG setup. B: Oddball task diagram. C: Experiment location sample.

Consistent with the methodology from our workgroup (Kuziek et al., 2017; Robles et al., 2021) the task was delivered via a Raspberry Pi 2 Model B computer using Version 9 of the Raspbian Stretch operating system and version 2.7.13 of Python. The Raspberry Pi was powered using a Microsoft Surface Pro 3 laptop. The task tones were delivered using Sony earbuds connected to a standard 3.5mm audio connector in the Pi computer. Using the GPIO pins in the Raspberry Pi 2, 8-bit TTL pulses were sent to the EEG amplifier through a wired connection.

The experiment was carried out in three different bicycle lanes nearby the north campus of the University of Alberta. During the time of data collection (August-October 2019), such lanes were characterized by different levels of traffic: (heavy traffic, intermediate traffic, low traffic). Figure 1C shows a sample location of one of the experimental conditions. The heavy traffic condition (figure 2A) is located at the Saskatchewan Drive road lane (https://goo.gl/maps/BQG8ERJoEzcajz2WA). This lane is characterized as a multi-use path with a flow of fast traffic (50 km/hr) alongside the cycling space. Therefore, the ongoing sounds on the fast traffic should interfere with the participant’s cognitive resources during the oddball-cycling task. This level of cognitive interference can be expected to be higher in heavy traffic and lower in quiet traffic. This lane is located in a designated multi-use path and cyclists are protected from car traffic.

**Figure 2.**
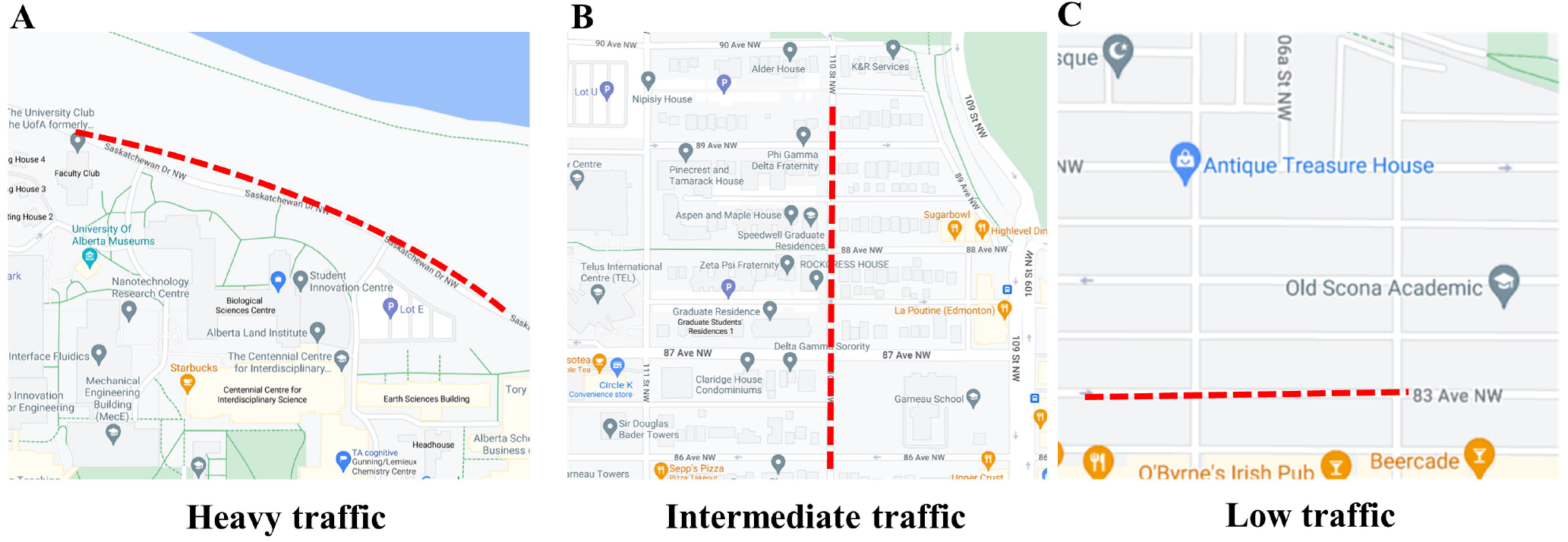
Experimental Conditions. A: Heavy traffic. B: Intermediate traffic. C: Low traffic.

The next condition is the moderate traffic condition (Figure 2B). This lane was a painted bike lane located on 110 Street NW (https://goo.gl/maps/7v6y3UJUG3kPVm6v6). The traffic volume in this lane is lower than the heavy condition as it is located in a smaller, one-way street of local traffic. Importantly, this is a non-protected, striped bike lane. Therefore, since drivers pass close to the cyclists (> 2 meters), the sound of the passing cars is expected to interfere with the oddball task to a lesser degree than in heavy traffic. The low-traffic condition (Figure 2C), located at 83 Avenue NW (https://goo.gl/maps/RmKJCX97od5ARQE97), is a new protected-buffered bike lane characterized by the lowest traffic among all lanes. Given the characteristics of this lane, one can expect that cyclists will experience the least level of traffic noise interference in this condition. During the experimental conditions, participant speed/safety was enforced by a research assistant who rode in front of the participant at an average distance of 12 feet. The assistant was trained to slow down and come to a full stop if necessary (e.g., incoming large vehicles at intersections, etc.).

Participants completed an auditory oddball task (Squires et al., 1975). In this task, participants listened to a series of tones played consistently via headphones (1,000-Hz “standard” or 1,500-Hz “target” tone played at 65 dB). Participants were instructed to use the button to respond to the target tone and to withhold a button response to the standards. Participants responded to each target tone during a delay period (1,000-1,500 ms) that followed after each tone presentation during the experiment. Each experimental condition contained a set of 250 trials with a total of 20% of these being the target tone (Figure 1B). Each condition lasted approximately 6 minutes. At the end of each condition, participants and research assistants rode the bicycles to the starting point of the following condition. Participants were given a small break between experimental conditions during which impedance levels were checked. The order of the conditions was randomized among participants.

### 2.3. EEG Recording

The EEG signal was collected using active, wet, low-impedance electrodes (actiCAP active electrodes kept below 10 kΩ). At the start of the experiments and between conditions, impedance was lowered using a blunted syringe with SuperVisc electrolyte gel. Streamed data were visually inspected to ensure low levels of impedance. Using the 10-20 electrode system, we employed the following electrode set: F3, F4, T7, T8, C3, C4, P7, P8, P3, P4, O1, O2, Fz, Cz, Pz. Location AFz was used for the ground electrode. Ag/AgCl disk electrodes. Data were recorded online, referenced to the left mastoid, and re-referenced to the arithmetically derived average of the left and right mastoids offline.

A V-Amp 16-channel amplifier (Brain Products GmbH) was used to record the EEG data. The V-Amp was connected to the Microsoft Surface laptop where the EEG was recorded using Brainvision Recorder software (Brain Products GmbH). Additionally, passive Ag/AgCl easy cap disk electrodes were used to record the vertical and horizontal bipolar electrooculography (EOG) via Bip2Aux adapters. EOG electrodes were affixed vertically above and below the left eye and affixed horizontally 1 cm lateral from the outer canthus of each eye. Nuprep exfoliating cleansing gel (Weaver & Co) was used to clean the participant’s skin before electrode placement. As previously mentioned, electrolyte gel was applied to each electrode to maintain inter-electrode impedance under 10 kΩ. EEG Data was digitized at 1,000 Hz with a resolution of 25 bits and hardware filtered online between 0.1 and 30 Hz, with a time constant of 1.5155 and notch filter at 60 Hz. All the experimental equipment was carefully placed inside a Lululemon backpack. The Raspberry Pi, connection cables, headphones, and electrodes were strategically placed inside the backpack pockets and secured with Velcro bands to reduce cable sway and tension that could disconnect signals during data collection. The overall weight of the backpack was 4.55 lbs.

### 2.4. EEG Processing

Analyses were computed in Matlab R2019a (MathWorks) using EEGLAB (Delorme & Makeig, 2004) and custom scripts (https://github.com/kylemath/MathewsonMatlabTools). Statistical analyses were computed on JASP (JASPTeam, Amsterdam, Netherlands). The EEG markers were used to construct 1200-ms epochs (200 ms pre-stimulus baseline) time-locked to the onset of standard and target tones, with the average voltage in the first 200-ms baseline period subtracted from the data for each electrode and trial. Variance in the data due to horizontal and vertical eye movements, as well as eye blinks were regressed out using the Eye Movement Correction Procedure (Gratton et al., 1983). An artifact rejection was applied and any trials with voltage differences from baseline larger than +/− 500 μV were removed. The resulting number of rejected trials following procedures did not vary significantly across experimental conditions (*F*(1.54,35.43) = 0.120, *p* = .83).

### 2.5. ERP Analysis

For the N1 and P2 analyses, time window cutoffs were chosen based on the previous EEG cycling studies from our workgroup (Scanlon et al., 2020). The N1 time window was defined as the grand-average of the negative increase in amplitude between 118 and 218 ms, maximal at electrode Fz. This component showed a peak latency of 168 ms. The P2 time window was defined as the grand average of the positive increase in amplitude between 218 and 318 ms at electrodes Fz and Pz, with a latency peak of approximately 292 ms. The P3 time window was selected by computing a grand-averaged, difference-wave ERP across participants and conditions to avoid biasing the selected time window towards any condition). Using this grandaverage waveform, we selected the peak of positive increase in amplitude between 355 and 505 ms. We selected the highest peak within this time window, resulting in a peak latency of 430 ms.

We conducted one-way, repeated-measures ANOVAs to test for ERP differences between the different bicycle lanes in the electrodes of interest. For instance, to test for the N1 and P2 differences across bike lanes, an ANOVA was run using the bike lane factor (Sask. Dr., 110 St., 83 Ave.) for the standard and targets tones respectively. For the P3 analysis, we conducted a one-way, repeated-measures ANOVA using the target-standard difference-wave ERP between the different bicycle lanes. The significance criteria α was set to 0.05 for all comparisons. For the ERP and other ANOVAs in this manuscript, Sphericity corrections were applied when necessary.

### 2.6. Spectral analysis

In addition to the ERP analysis, we were interested in investigating differences in overall sustained alpha power between bike lanes. We conducted an EEG frequency spectra analysis on the averaged EEG epochs using the wavelet routine from the Better Oscillation Method, BOSC, (Hughes et al., 2012). The epochs consisted of a 1000 ms baseline and a 2000 ms post-stimulus period using the standard tones. We selected a 6-cycle wavelet transform across a frequency range of 0.1 Hz to 30 Hz, increasing in 0.5 Hz steps.

## 3. Results

### 3.1. Behavioral Results

Reaction times and accuracy were calculated for the three conditions. Figure 3A depicts the reaction times in participant response to the target oddball tone. For reaction time, using a Greenhouse-Geisser corrected one-way repeated measures ANOVA, we found a significant main effect for condition type (*F*(1.455, 33.46) = 3.90, *p* = 0.042, η^2^_p_ = 0.145). A Holm adjusted post-hoc test revealed a significant increase in reaction time in the low traffic condition lane relative to heavy traffic. (M_diff_ = −35.43, *p* = 0.01). No further significant differences in reaction times were found among the different lanes. Figure 3B depicts participant accuracy to the target tones in the oddball task. A one-way repeated measures ANOVA found no significant differences in accuracy among the conditions (*F*(2, 46) = 0.89, *p* = .415, η^2^_p_ = 0.038).

**Figure 3.**
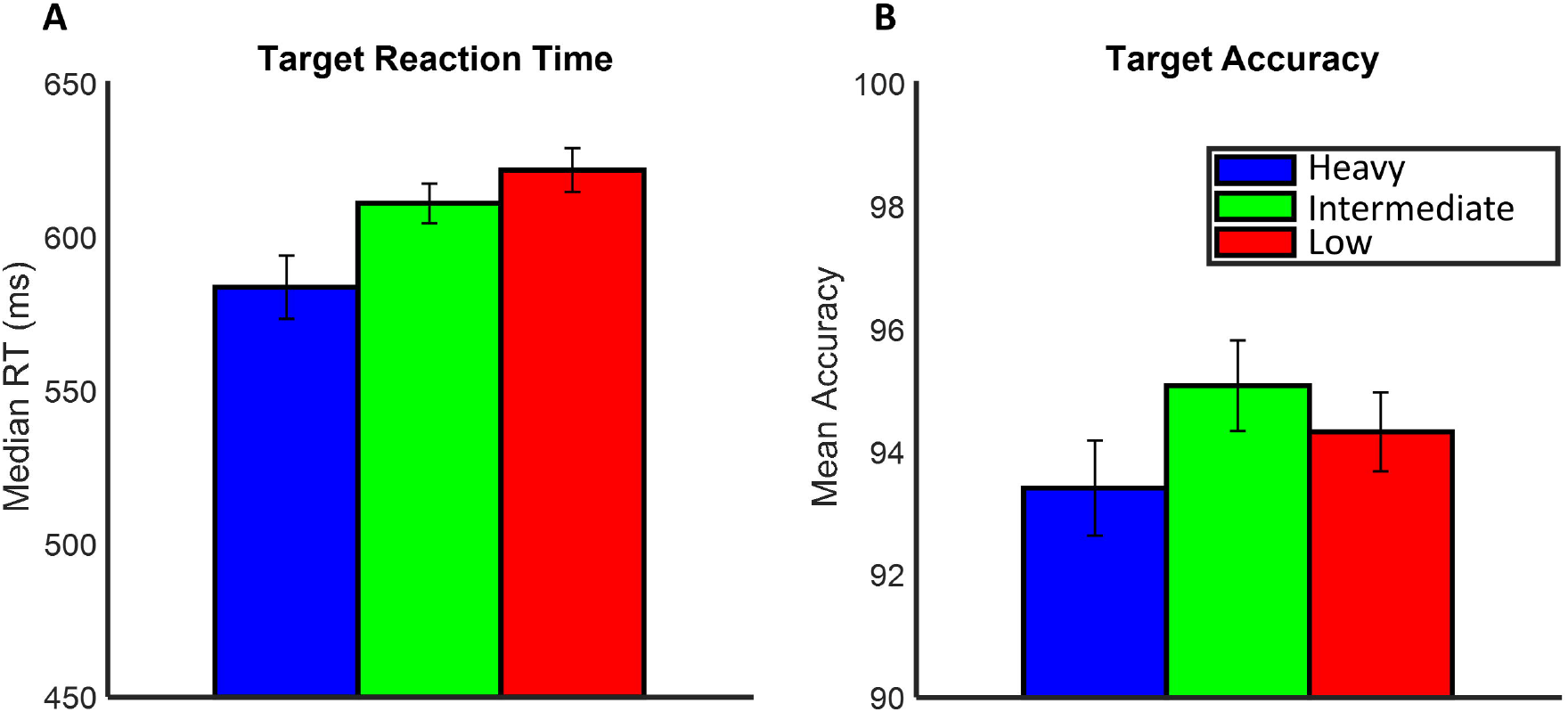
Behavioral results. A: Target accuracy. B: Target reaction time.

### 3.2. ERP Morphology and Topography

Figure 4 shows the grand average ERPs for standard and target tones at electrodes Fz and Pz for all traffic types. Standard tones are depicted in black while target tones are depicted in colored lines. The shared regions depict the standard errors at each ERP line. A visual inspection of the grand-average ERPs shows an overall increase in negative voltage for the N1 time window (118-218 ms), followed by a positive increase at the P2 time window (218-318 ms) at all electrodes and in all conditions. Furthermore, there is an increase in positive voltage at the P3 time window for the target tones relative to the target tones showing a maximal difference at Pz.

**Figure 4:**
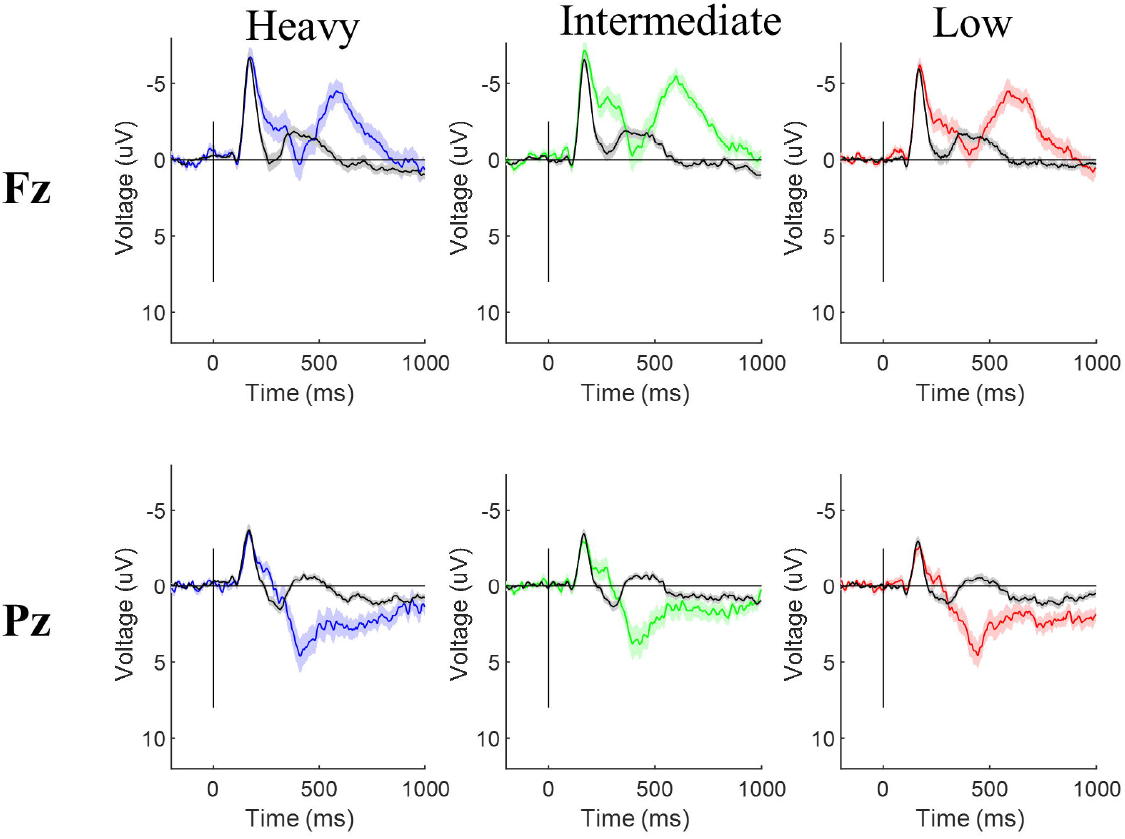
Grand-Average ERPs by traffic condition.

Figure 5A and 5B depict the topographies for the N1 and P2 time windows for the standard and target tones respectively. The topographies for the chosen N1 time window show a fronto-central distribution while for the P2 time window, the topographies show a posterior distribution where the highest activity is located towards the parietal region and seen more prominently in the standard tones.

**Figure 5:**
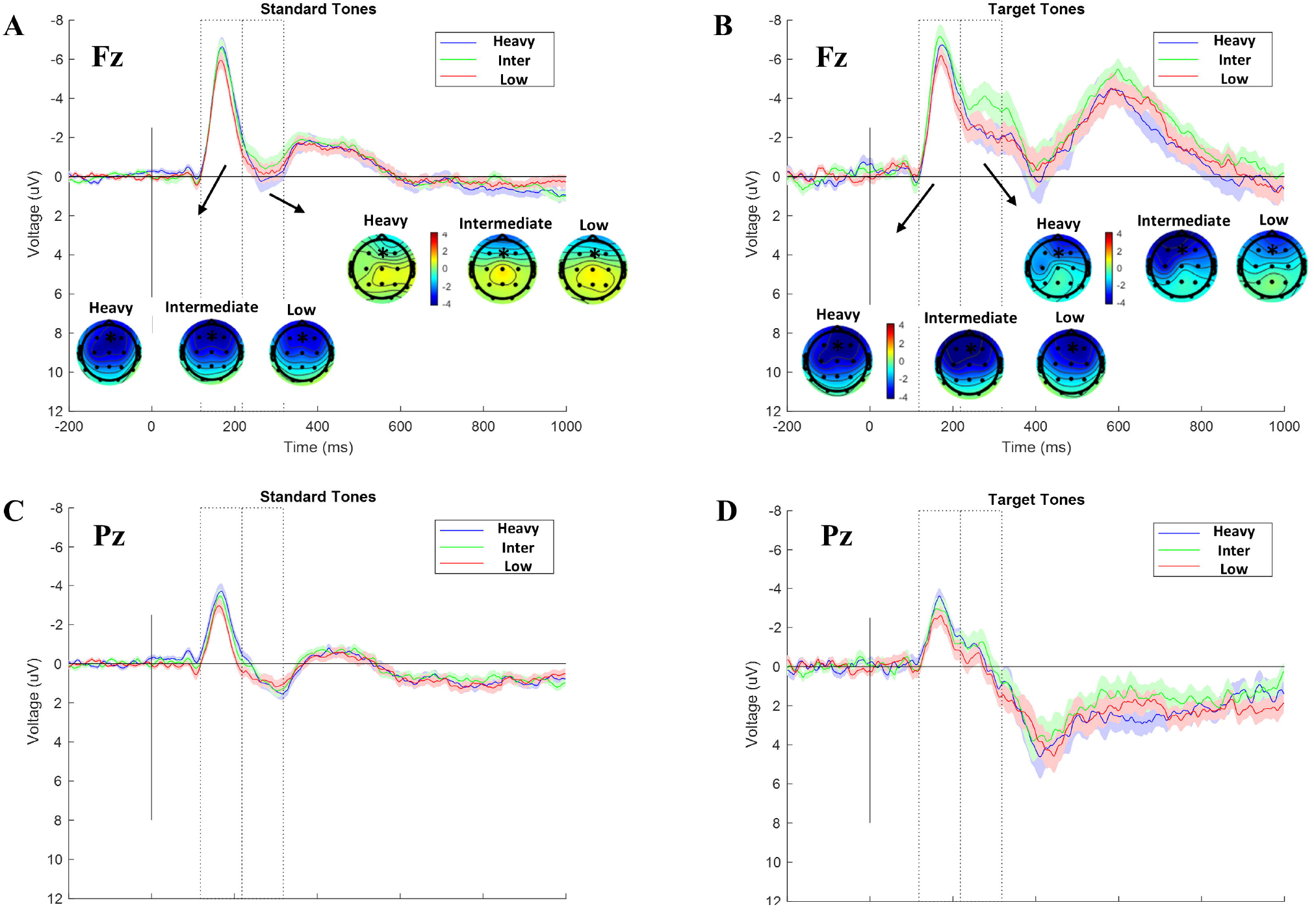
N1 and P2 ERPS. A: N1 and P2 ERP time windows for standard tones at electrode Fz. Topographies for the N1 & P2 by traffic condition for standard tones. B: N1 and P2 ERPs for target tones at electrode Fz. Topographies for the N1 & P2 by traffic condition for target tones. Shaded regions represent standard errors. C: N1 and P2 ERP time windows for standard tones at electrode Pz. D: N1 and P2 ERP time windows for target tones at electrode Pz. Shaded regions represent standard errors.

### 3.3. N1/P2 ERP results

We conducted a one-way repeated measures ANOVA to compare the mean in the N1/P2 ERP time window at electrodes Fz and Pz. Figure 5A shows the N1/P2 ERPs for standard tones at electrode Fz. We found a significant N1 main effect for the bike lane condition for standard tones (*F*(1.45, 33.517) = 3.63, *p* = 0.05, η^2^_p_ = 0.136). A Holm adjusted post-hoc test revealed a significant decrease in N1 amplitude in the low traffic condition relative to heavy traffic (M_diff_ = −0.61, *p* = 0.041). No significant differences in the P2 ERP were found for standard tones at electrode Fz (*F*(1.50, 43.63) = 1.49, *p* = 0.23, η^2^_p_ = 0.049).

Figure 5B shows the N1/P2 ERPs for target tones at electrode Fz. Regarding the N1 ERP, no differences in target tones were found between the bike lanes for electrode Fz, (*F*(2,46) = 2.42, *p* = 0.10). A repeated-measures ANOVA found a significant main effect for the traffic condition for the P2 time window for target tones (*F*(1.70,49.48) = 3.75, *p* = 0.037, η^2^_p_ = 0.115). A post-hoc revealed a less positive amplitude during the P2 time window for intermediate traffic relative to the low traffic condition (M_diff_ = −1.420, *p* = 0.032).

Figure 5C shows the N1/P2 ERPs for standard tones at electrode Pz. We found a significant main effect for the traffic condition for standard tones in the N1 time window (*F*(2, 46) = 7.42, *p* = 0.002, η^2^_p_ = 0.244). A Holm adjusted post-hoc test revealed a significant decrease in N1 amplitude in the low traffic relative to heavy traffic condition (M_diff_ = −0.82, *p* = 0.001). No statistically significant differences were found for the P2 ERP at electrode Pz for the standard tones (*F*(1.40, 40.81) = 0.87, *p* = 0.39, η^2^_p_ = 0.029). Figure 5D shows the N1/P2 ERPs for target tones at electrode Pz. No significant differences in ERP amplitude were found for the N1 (*p* = 0.34) or P2 (*p* = 0.44) ERPs for target tones.

### 3.4. P3 Results

Figure 7 shows the averaged difference-wave ERP for the target-standard tones at electrode Pz. The difference waveform shows an increase in positive power starting at approximately 300 ms that resembles the classic oddball-related P3 (Luck, 2014). A one-way repeated-measures ANOVA was conducted to test for differences in P3 amplitude between the bicycle lanes. The ANOVA found no statistically significant differences in P3 amplitude between the traffic conditions (*F*(2,46) = 0.191, *p* = 0.82).

**Figure 6:**
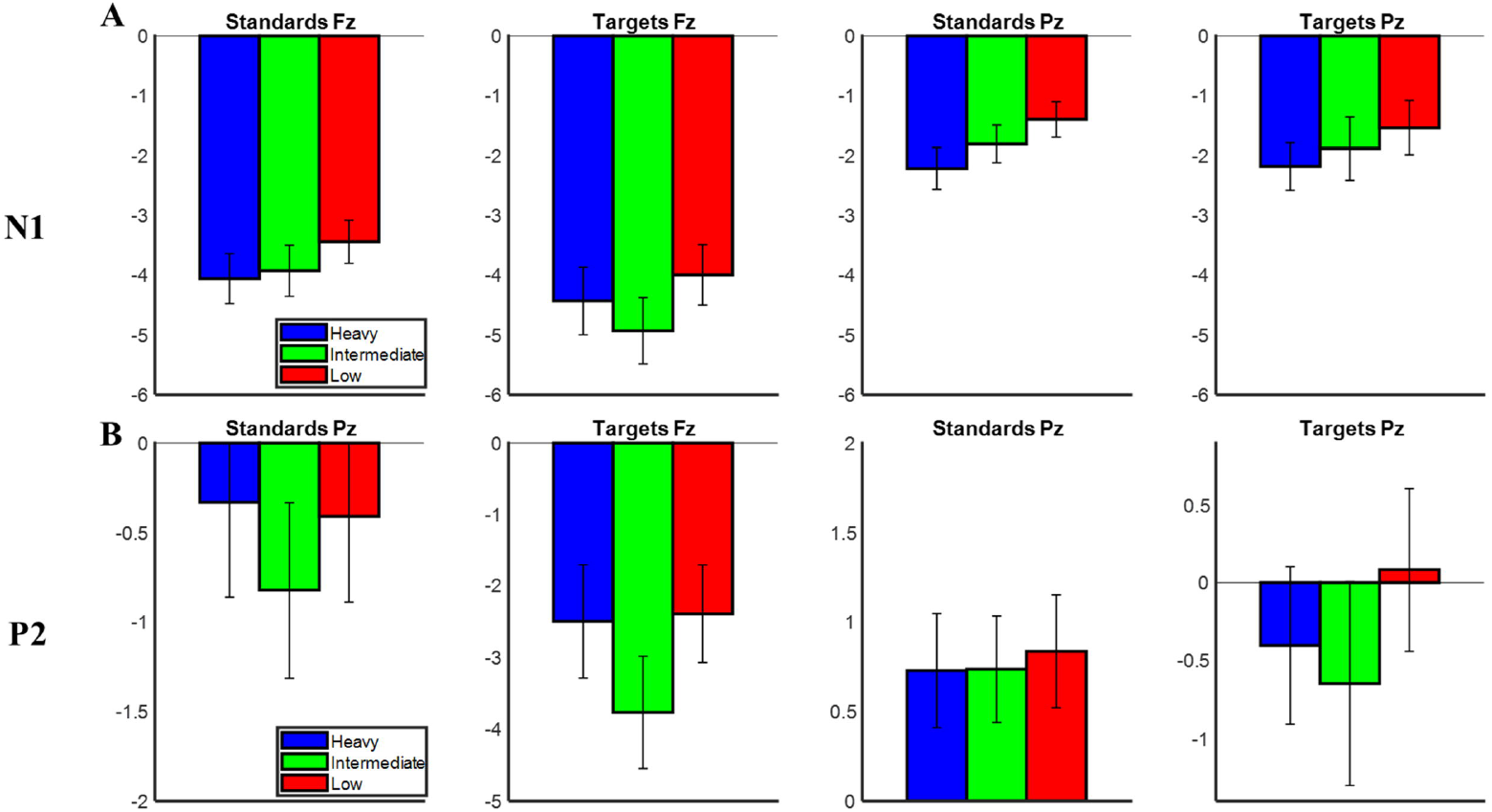
Bar graph summaries of the N1 and P2 ERP comparisons by electrode type and condition. A: N1. B: P2.

**Figure 7:**
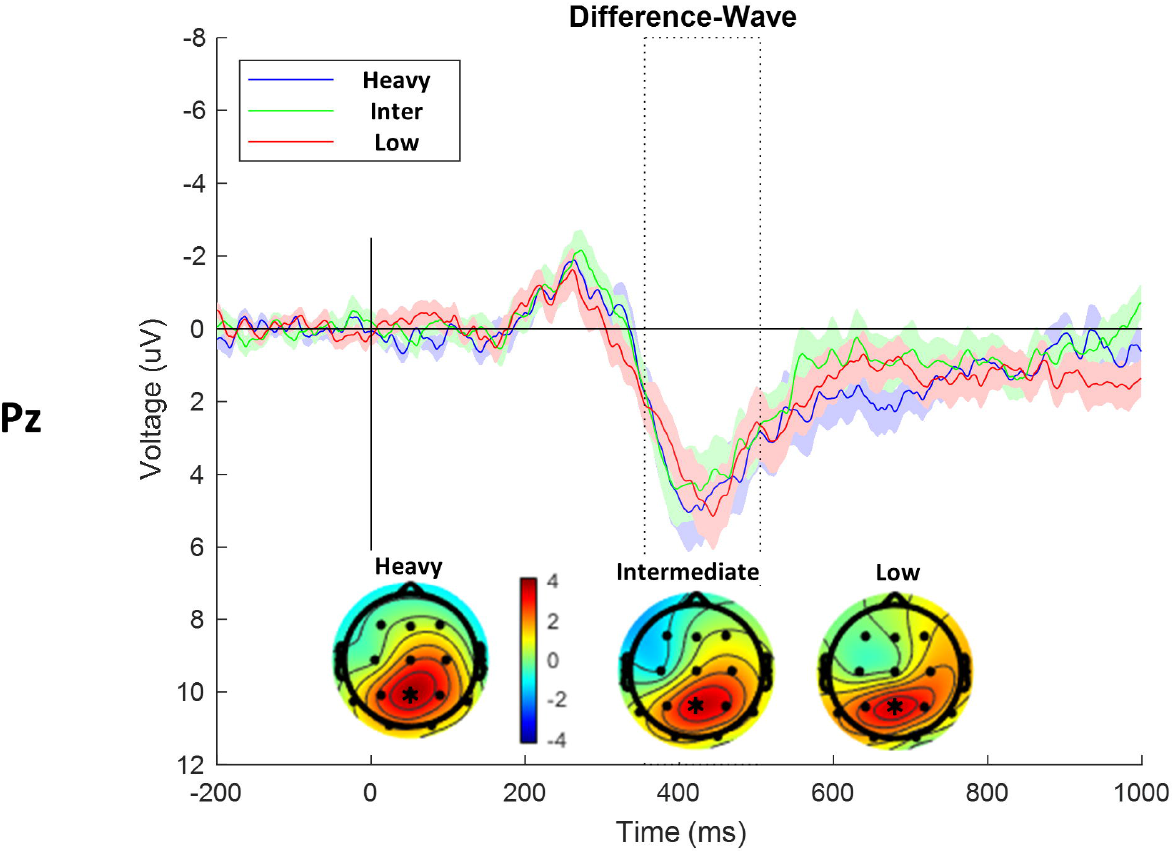
Difference-wave P3 ERPs at electrode Pz. Topographies for the P3 time window. Shaded regions represent standard errors.

### 3.5. Spectral Results

Figure 8 shows the EEG power spectra plots for the different lanes. The different color lines represent the different bike lanes. The shaded regions depict the standard error among each group. The 1/f EEG spectra plots show a slight increase in power within the alpha range (8-12 Hz) for all conditions. It also depicts the topographies in the alpha range by experimental conditions which exhibit similar distributions in the parietal regions. Notably, the figure shows a decrease in alpha power in the low traffic condition relative to the other traffic conditions. We conducted a one-way repeated-measures ANOVA to test for differences in alpha power between the road types. The ANOVA revealed a significant main effect for bike lane at electrode Pz (*F*(2, 46) = 3.98, *p* = 0.02, η^2^_p_ = 0.148). A Holm-corrected post-hoc comparison shows a statistically significant difference in alpha power in the low traffic condition relative to the intermediate traffic (*p* = 0.021).

**Figure 8:**
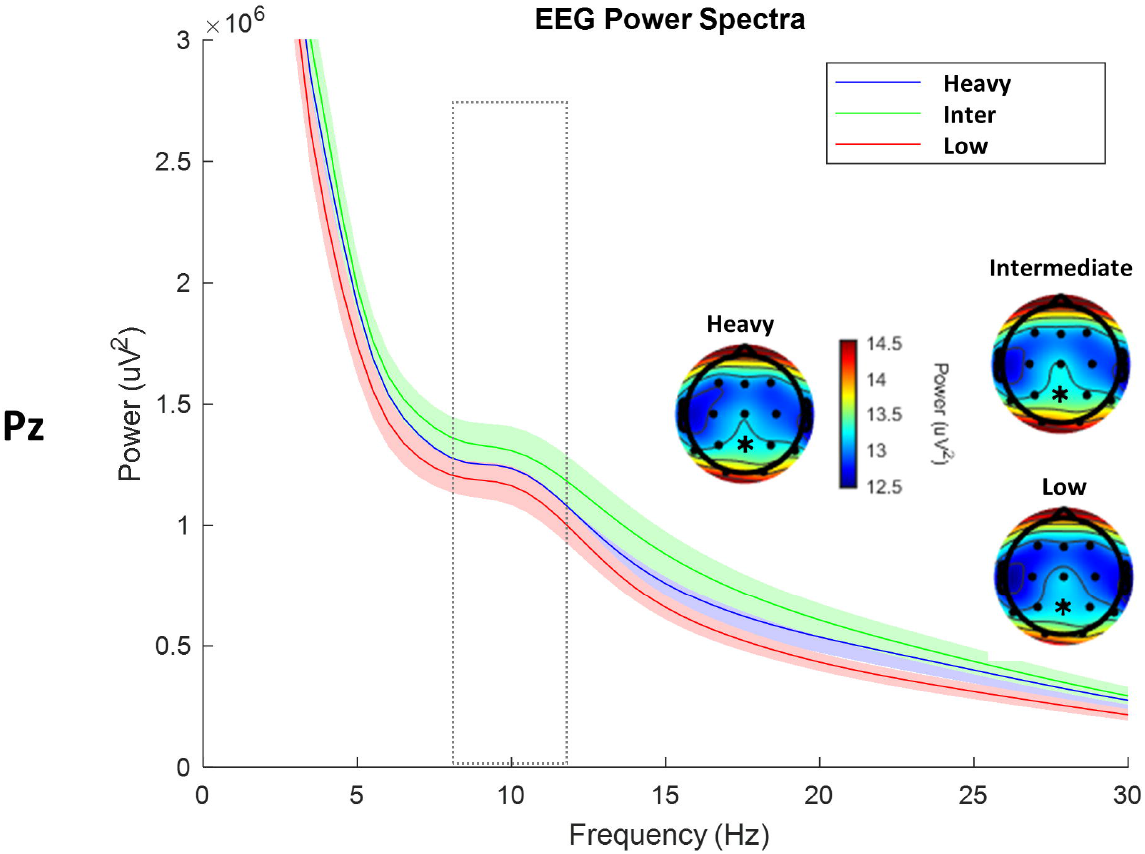
EEG power spectra at electrode Pz. The squared area depicts the alpha range of 8-12 Hz. Shaded regions represent standard errors.

## 4. Discussion

In this study, we measured EEG activity from individuals under different urban traffic conditions. Participants completed an auditory oddball task while riding a bicycle in three different cycling environments. The first condition (low traffic) was recorded in a separated cycling lane located next to a one-way, low traffic road. The second condition involved completing the oddball task alongside a painted, intermediate traffic volume lane, and the final condition involved riding on a multi-use path alongside heavy/fast traffic. The task parameters such as the riding speed, stimulus presentation, and recording methods were chosen following previous mobile EEG work from our laboratory to ensure our methodology is replicable under new environments and for comparability. Overall, we found evidence suggesting that changes in urban traffic environments can be reflected in auditory mechanisms, such as the N1 increase under heavy traffic.

### 4.1. Behavioral Results

In the present paradigm, participants responded to the target tones in the oddball task by pressing a button attached to the handlebar. We calculated button response accuracy and response times in each of the experimental conditions. We found no significant differences in accuracy by traffic condition. While there is a non-significant difference between accuracy scores, there was slightly higher accuracy in the low traffic condition relative to the heavy traffic condition. However, participants performed better during the intermediate traffic condition overall. We found significant statistical differences in target reaction time between the traffic conditions. Interestingly, participant reaction time was statistically slower in the low traffic lane and faster under heavy traffic. While it is generally expected that fewer distractions lead to better performance (e.g. lower reaction time), it might be plausible that the difference in reaction time might be influenced by participant arousal levels. Given the linear relationship between reaction time and traffic volume, the faster/heavy traffic condition might have increased participants’ subjective arousal, thus leading to faster button response (Kovacs & Bories, 2010; Nishisato, 2013). In other words, participants’ reaction times might have slowed down under low traffic due to a more relaxed state as opposed to riding alongside heavy traffic. Lastly, while not reported here, the proportion of targets responded to in the heavy traffic was lower than the other traffic conditions. While not statistically significant, this decrease in response might suggest that higher attentional demands during heavy traffic can affect task performance.

### 4.2. N1 and P2 differences by traffic volume

In this study we hypothesized that, relative to low traffic, heavy traffic would elicit a more negative N1 amplitude during street cycling. Figure 6 shows a summary of the comparisons performed for this analysis. Consistent with previous work from our laboratory, the main finding in the current study is that increases in environmental noise were associated with increases in N1 amplitude (see Scanlon et al., 2017; Scanlon, Townsend, et al., 2019). Namely, we found that relative to low traffic, heavy traffic was associated with a significant increase in N1 to the standard tones at both electrodes Fz and Pz. The significant differences in N1 amplitude found in the present study occurred between the heavy vs low traffic only. Interestingly, no significant differences were found between the intermediate and low traffic conditions. However, a visual inspection of the N1 shows a consistent pattern where the intermediate traffic amplitude falls between high traffic and low traffic (Figure 6A). In other words, as the traffic volume goes from heavy > intermediate > low, the observed N1 amplitude follows the same order: higher amplitude in heavy traffic, medium amplitude in the intermediate traffic, and lowest amplitude in low traffic. Due to the randomization of task orders in this study, we attribute these differences to environmental conditions and not order effects.

The role of the auditory N1 in stimulus processing has been established in previous literature. It is known that changes in the properties of auditory stimuli (e.g., intensity and frequency) modulate N1 amplitude (Näätänen & Picton, 1987). For instance, changes in the environment are followed by changes in N1 activity (Paiva et al., 2016). In the context of mobile EEG, our workgroup’s focus on the N1 started in a study by Scanlon et al. (2017). Using an oddball task, the recorded subject’s EEG while sitting indoors and while riding a bicycle outside. They found an unexpected increase in N1 and a decrease of P2 during outdoor cycling relative to sitting indoors. In a follow-up control experiment, we found that this N1/P2 effect was larger when subjects were listening to outdoor sounds and white noise relative to noises of silence. A follow-up experiment further showed that increases in traffic sounds, as indexed by the road type (quiet park vs noisy road) further showed the previously found N1 effect (Scanlon et al., 2020). In these studies, the authors concluded that the observed modulation in N1 could serve as a filtering mechanism that should facilitate sound processing in louder environments.

In the present study, we were interested in exploring the influence of environmental conditions during active bicycle riding. By deploying our paradigm to different urban settings, we tested whether it was possible to obtain reliable N1 effects given the differences in traffic levels. The current results show that the N1 effect shown by our workgroup in parks vs roads is also sensitive enough to quantify differences in auditory processing in cycling environments. In other words, we showed that even in similar urban environments, differences in traffic conditions can influence auditory processes during the oddball task and while performing a motor task. These findings can inform city infrastructure and planning by showing how the volume of vehicles interacts with attentional processes in cycling, and by providing a more objective measure of traffic noise impact on bike infrastructure.

Given the mixture of behavioral results and the consistency of the N1 effect, one must consider the behavioral implications of the observed N1 differences of the present study. Like in Scanlon et al., (2019, 2020), we did not find significant differences in target accuracy. This could lead to the conclusion that a larger sample size might be necessary to better quantify any statistical differences in target accuracy. But as noted in the behavioral discussion, we observed a trend in response accuracy where participants responded more accurately to the oddball task in the low traffic relative to heavy traffic. However, the highest accuracy was found in the intermediate traffic conditions. Of equal interest is the difference in target reaction times between low and heavy traffic. We found slower reaction times in the low traffic condition and faster reaction times in heavy traffic. An important observation is that the N1 effect was not found in the target conditions. Whether the lack of significant target N1 differences is attributed to the sample size, ultimately, the present study cannot confirm whether the N1 effect is associated with behavioral accuracy as previously found (Hink et al., 1977).

We only found one significant difference in P2 amplitude between conditions, a less positive amplitude in the intermediate traffic relative to low traffic. Given the previous findings from our group showing a decrease in P2 amplitude during outdoors relative to indoors (Scanlon et al., 2019), and a similar decrease in outdoor sounds vs silence noise (Scanlon et al., 2019), we expected to find a decrease in P2 amplitude in the heavy traffic relative to low traffic. The only significant P2 difference found in the present study was between intermediate and low traffic, where there was a decreased P2 amplitude in the intermediate condition. Interestingly, this effect was only significant in the target tones. While inconsistent with our initial expectations, the lack of P2 differences among the other comparisons might suggest a general role in P2, where even similar road conditions (regardless of noise intensity) can elicit an equal decrease in this ERP (Scanlon et al., 2020). An alternative explanation to these findings has to do with the consistency of traffic sounds. Because heavy traffic is characterized by consistently loud noises and low traffic by consistently quieter noises, it is possible that in the intermediate condition participants were exposed to a switch between noisy and quiet. This would make the auditory filtering associated with the P2 harder to predict. In the context of other mobile research, other studies have observed decreases in P2 amplitude during increased task load (Reiser et al., 2021). Given the fact that P2 amplitude is generally associated with task difficulty (Sugimoto & Katayama, 2013), an important question for future mobile paradigms is to test whether the P2 effect reflects a filtering function that discriminates between outdoor and quiet sounds or whether increases in bike difficulty can elicit changes in P2 amplitude.

### 4.3. P3 differences by traffic volume

Regarding the P3, it was hypothesized that riding in heavy traffic would be associated with decreases in P3 positivity. Given the well-established literature associating P3 amplitude to task effort (Kok, 2001; Polich & Comerchero, 2003), we expected to test whether changes in traffic conditions alongside the bike lanes would be reflected in changes in P3 amplitude (e.g., a more negative amplitude during heavy traffic). There were no significant differences in P3 amplitude between the experimental conditions. This lack of a P3 difference using a free navigation paradigm brings several points to discuss when it comes to studying attentional demands using motion paradigms. First, the P3 effect is generally associated with task effort (Dinteren et al., 2014) and has been historically studied in stationary indoor laboratory settings. The present findings are consistent with a previous skateboarding study where we found that increases in task difficulty were not associated with changes in the P3 (Robles et al., 2021). A common factor between the current findings and the skateboard findings is that we compared several experimental conditions involving free navigation and with similar motor and sensory demands. For example, in this study, the riding conditions did not vary in terms of motor intensity (terrain incline and riding speed remained constant among the bicycle lanes), and all the conditions were carried out in unstructured outdoor environments where participants are exposed to outdoor stimuli. In terms of task interference, the lack of a P3 effect in our studies supports the findings of several mobile studies showing that free unconstrained navigation appears to interact with attentional resources differently from stationary conditions and other conditions that involve fixed movements. Some of these studies are discussed below.

Scanlon and colleagues (2017) found no differences in P3 amplitude between stationary pedaling and sitting on a stationary bike while participants completed an oddball task in an EEG chamber. These results suggest that an increase in motor demands alone did not modulate P3 amplitude. However, in a follow-up study, Scanlon and colleagues (2019) showed that relative to an indoor sitting condition, riding outdoors was associated with a significant P3 decrease. In a cycling study that also used an oddball paradigm, Zink and colleagues (2016) found an increase in P3 amplitude in a free condition relative to a fixed bike. The authors concluded that, in the free biking conditions, participants were exposed to a greater cognitive load relative to fixed cycling. Notably, Ladouce et al. (2019) used a series of oddball experiments to show that relative to standing still, walking is associated with a decrease in P3 amplitude. Crucially, they also found similar P3 amplitudes decreases between walking and when participants were pushed while sitting in a wheelchair. The authors also found similar P3 amplitudes between treadmill walking and standing still, arguing that the combined effect from visual and inertial stimulation was a likely candidate to explain the findings in their experiment. Taken together, the findings in the present study and the mobile studies discussed above might suggest a stronger attenuation of P3 amplitude in paradigms when we consider the transition from stationary and fixed to free movement. This is particularly true in outdoor studies where one can expect an increase in stimulus processing relative to traditional laboratory settings (Scanlon et al., 2019). Free navigation and outdoor paradigms are promising research alternatives relative to traditional indoor research (Makeig et al., 2009). Researchers must continue to explore the role of attentional processes using naturalistic paradigms to better understand human attention under higher ecological validity.

### 4.4. Alpha power differences by traffic volume

We hypothesized that, relative to low traffic, riding under heavy traffic would decrease alpha power during the oddball task. We found no differences in alpha power between the low and high traffic conditions. Interestingly, the only two conditions that yielded statistically different levels of alpha power were the low and intermediate, with a larger alpha power decrease in the low traffic. In a previous study (Robles et al., 2021), we were motivated to test alpha power during skateboarding because previous mobile research has shown that increases in cognitive and motor load during motion are associated with decreases in power (Storzer et al., 2016; Zink et al., 2016). We showed that relative to a resting baseline, alpha power was greatly diminished during skateboarding. In that study, we found no differences in alpha power during increased cognitive and motor inference. We concluded that the difficulty of the task (completing an oddball task while riding the skateboard) would draw enough cognitive resources to deplete alpha power during the task. In terms of the present study, it is also possible that the task (riding a bicycle outdoors) led to a global state of cortical excitation due to the external environment stimulation and task. In this sense, the constant exposure to external stimuli during the three experimental conditions might have resulted in similar levels of cortical excitation. This increased excitation might lead to equal decreases in alpha amplitude in this study (Sauseng et al., 2009).

### 4.5 Limitations and future directions

There are several limitations in this study that will be discussed in this section. We failed to obtain an objective measure of traffic volume of the road conditions during the time of the experiment (pre-pandemic traffic). We could have correlated the N1 amplitude to the levels of traffic noise in dB during the time of the experiment. Future mobile experiments testing auditory paradigms outdoors can benefit from recording and quantifying the levels of street noise. For example, in tasks that involve auditory filtering, recording the outdoor noise levels can allow for the correlation analyses of ERP or spectral power to the decibels of the street sounds. We also did not record a baseline condition with no traffic sound. Given our interest in testing differences in N1 in similar urban environments, we did not consider recording a condition without traffic sounds. Implementing a condition where participants ride in a busy environment (busy track or park) could have provided an important piece of information regarding the role of the N1. For instance, a busy park without traffic could serve as a suitable baseline condition where subjects are equally busy as in an urban track but without traffic sounds. In a previous cycling study, Scanlon et al. (2020) showed marked differences in N1 amplitude between a quiet park and near a roadway. To further assess the degree to which the reported N1 effect is sensitive to environmental sounds it could be helpful to integrate a condition equally busy as a street lane but without traffic.

Due to the real world nature of our experiment, our three conditions had other differences between them besides just the amount of traffic. The faced different directions, had difference scenery, had different numbers of intersections, different lighting and trees, and different amount of other bike traffic, and these variables changed from day to day outside of our control. These degrees of freedom are an inevitable consequence of this realistic scenario research. We find it easier to consider the experimental manipulation as including all these variables together as a package, but this does make replication difficult. Further difficulties are present due to changing infrastructure. Our intermediate style pathway has since been replaced with a fully separated bike lane more similar to the low traffic lane since the time of data recording. Further work might use virtual reality or full controlled outdoor scenarios to better control these extraneous variables.

An important consideration is how much we can manipulate the task difficulty in a real-world setting without compromising participant safety. While the lack of behavioral effects in the study can be considered a limitation, there is a limit to which we can increase task difficulty before it can turn into a falling hazard. For instance, using flashing LED glasses, we have tested a visual version of the current study in an indoor running track while subjects ride a Bluetooth-operated skateboard. A disadvantage of this design is that it can only be deployed to semi-controlled settings (e.g., using an empty running track) where any accidents can be minimized. For example, in the current study, we had a researcher riding in front of the participant to ensure participant safety at all times. While we had no forms of safety-related incidents or situations, it is possible that attending to a visual task might affect participant’s coordination, which is an undesirable outcome as we prioritize participant and equipment safety.

Another consideration for future designs is using a webcam to time and capture moments from the environment. In previous mobile studies, researchers have implemented the use of recording environmental events during driving such as pedestrian crossing and cars driving nearby the participants (Di Flumeri et al., 2018). Other indices of attention, such as eye blinks, are now being implemented to record cognitive engagement during navigation (Wunderlich & Gramann, 2020). These events could be used for interesting analyses (such as evaluating EEG signals and reaction time at random times such as during a change of traffic light or a sudden distractor). Furthermore, using a structured protocol to record outside conditions (weather conditions, infrastructure condition, average environmental noise, average level of stops/nearby distractions, etc.) could provide rich details that could be used in conjunction with task performance and EEG recordings.

## Conclusion

In the current study, we used an oddball EEG cycling paradigm to test the effects of environmental noise in auditory attention while participants rode in different urban lanes. We showed that levels of traffic volume were associated with changes in auditory attention. While showing the feasibility of deploying mobile paradigms into real-world environments, we demonstrated that traffic volume interacts with cognitive resources during multitasking.

## Notes

https://github.com/APPLabUofA/Bike2

### Competing Interest Statement

The authors have declared no competing interest.

